# The tempo of trophic evolution in small-bodied primates

**DOI:** 10.1101/2020.03.17.996207

**Authors:** Jeremiah E. Scott

**Affiliations:** Department of Medical Anatomical Sciences College of Osteopathic Medicine of the Pacific Western University of Health Sciences 309 East Second Street Pomona, California 91766 U.S.A.

**Keywords:** body size, diet, insectivory, herbivory, heterotachy

## Abstract

**Objectives:** As a primary trophic strategy, insectivory is uncommon and unevenly distributed across extant primates. This pattern is partly a function of the challenges that insectivory poses for large-bodied primates. In this study, I demonstrate that the uneven distribution is also a consequence of variation in the rate of trophic evolution among small-bodied lineages.

**Methods:** The sample consisted of 307 species classified by primary trophic strategy and body size, creating an ordered three-state character: small-insectivorous, small-herbivorous, and large-herbivorous. I tested for rate heterogeneity by partitioning major clades from the rest of the primate tree and estimating separate rates of transition between herbivory and insectivory for small-bodied lineages in each partition.

**Results:** Bayesian analysis of rate estimates indicates that a model with two rates of trophic evolution provides the best fit to the data. According to the model, lorisiforms have a trophic rate that is 4–6 times higher than the rate for other small-bodied lineages.

**Conclusions:** The rate heterogeneity detected here suggests that lorisiforms are characterized by traits that give them greater trophic flexibility than other primates. Previous discussions of trophic evolution in small-bodied primates focused on the low frequency of insectivory among anthropoids and the possibility that diurnality makes insectivory unlikely to evolve or persist. The present study challenges this idea by showing that a common transition rate can explain the distribution of insectivory in small-bodied anthropoids and nocturnal lemurs and tarsiers. The results of this study offer important clues for reconstructing trophic evolution in early primates.

## 1 INTRODUCTION

Primates exhibit an impressive diversity of trophic strategies. Among extant members of the order, frugivory is the most widespread primary strategy, but folivory is also common (Gómez & Verdú, 2012; Kay & Covert, 1984). The clade also includes specialized lineages such as the graminivorous gelada (*Theropithecus gelada*) of the Ethiopian Highlands, the tree-gouging, exudativorous marmosets (genera *Callithrix*, *Mico*, *Callibella*, and *Cebuella*) of Amazonia and the Atlantic Forest, and the exclusively faunivorous tarsiers (family Tarsiidae) of the Malay Archipelago (Fleagle, 2013). Explaining how this diversity arose—and particularly how it has been shaped by other aspects of primate biology—is a major goal of evolutionary primatology.

Body size has been recognized as an important influence on primate trophic evolution since Kay (1975) noted that folivores are mostly large-bodied whereas insectivores are mostly small. The correlation between body size and diet has been attributed to two other size-related trends (Kay, 1975; Kay & Covert, 1984; Kay & Hylander, 1978; Kay & Simons, 1980). First, because insects are small, dispersed, and often elusive, acquiring enough of them to meet metabolic requirements becomes more challenging as body size increases and is probably physiologically impossible above a certain threshold without specializing on social insects (McNab, 1984). Second, as body size decreases, digestive retention time becomes shorter and metabolic rate per unit mass increases, making it difficult for small-bodied primates to extract sufficient nutrition from leaves, which are resistant to chemical digestion and must be slowly fermented in the gut (Lambert, 1998). These arguments have also been invoked to explain why large-bodied frugivorous primates rely on leaves as their main source of dietary protein whereas small-bodied frugivores are dependent on insects (Kay & Simons, 1980; Kay & Covert, 1984). Size differentiation between herbivores and faunivores is pervasive across mammals (Price & Hopkins, 2015; Grossnickle, 2020), indicating that the pattern found in primates is a general feature of mammalian biology.

The distribution of trophic strategies within small-bodied primates has generated additional hypotheses of constraint on the evolution of insectivory. Although insects are an important resource for many diurnal primates (e.g., Digby, Ferrari, & Saltzman, 2007; Kinzey, 1992; Souza-Alves, Fontes, Chagas, & Ferrari, 2011; Zimbler-DeLorenzo & Stone, 2011), insectivory as a primary trophic strategy (i.e., at least 50% a species’ diet) is found mainly in nocturnal lineages (Figure 1). The reason for the rarity of insectivory in small-bodied diurnal lineages is unclear, but one hypothesis that has been proposed is that competition with diurnal birds has limited the ability of primates to become established in the diurnal arboreal insectivore niche (Cartmill, 1980; Charles-Dominique, 1975; Ross, 1996). This idea is difficult to test, but direct interactions between the two clades certainly do occur (Heymann & Hsia, 2015), and there is evidence that such interactions have had an influence on the distribution of species in each clade (Beaudrot et al., 2013a, 2013b).

**FIGURE 1.**
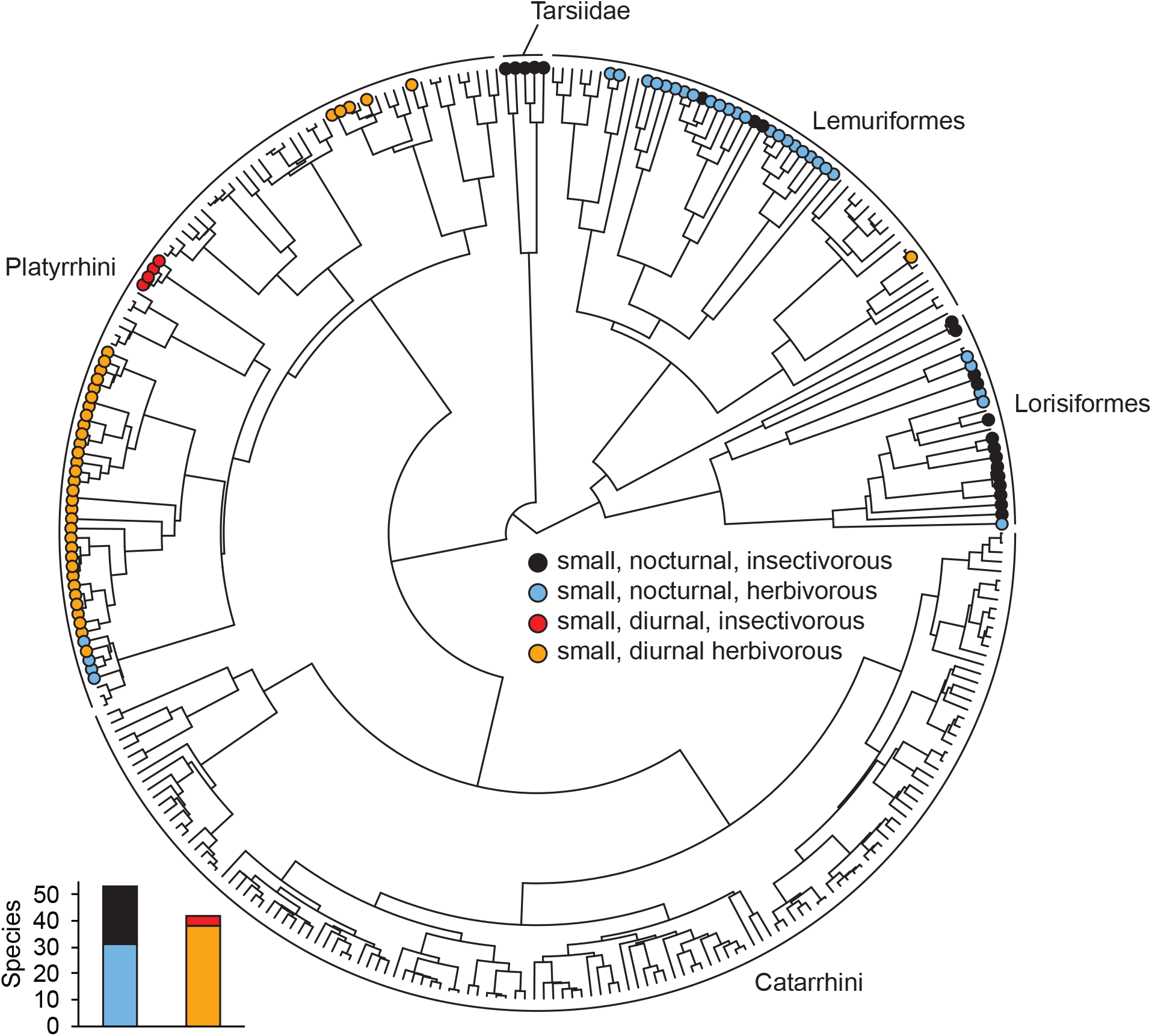
Primate phylogenetic tree from Springer et al. (2012) showing the distribution of trophic states by activity pattern among small-bodied species (<1 kg) at the tips (Scott, 2019). Cathemeral species are grouped with diurnal species. Large-bodied species, which are uniformly herbivorous and mostly diurnal, are not labeled. The relative frequency of insectivorous species is much greater among nocturnal lineages (41.5%) than among diurnal lineages (9.5%).

A long history of adaptation to herbivory has also been identified as a possible constraint. In his discussion of primate origins, Rosenberger (2013) advocated for the idea that frugivory was the formative trophic influence on early primate evolution, responsible for many of the apomorphies that distinguish primates from other mammals (Sussman, 1991; Sussman, Rasmussen, & Raven, 2013; Szalay, 1968). As a consequence, he argued, a primarily insectivorous diet presents primates with “intense selective challenges” (Rosenberger, 2013, p. 886), making it difficult for them to switch from herbivory to insectivory. Studies of acidic mammalian chitinase genes (*CHIA*s) provide support for the idea that some primate lineages have experienced changes to their digestive biology that may decrease the likelihood of insectivory evolving or persisting (Emerling, Delsuc, & Nachman, 2018; Janiak, Chaney, & Tosi, 2018). However, in contrast to the global constraint proposed by Rosenberger, the distribution of *CHIA* pseudogenes and deletions among extant species indicates that gene functionality has been maintained by selection in some primate lineages and lost multiple times in others (Emerling et al., 2018; Janiak et al., 2018). As noted by the authors of the *CHIA* studies, this pattern of evolution is consistent with the hypothesis that insectivory was important for early primates (Cartmill, 1974, 1992, 2012), with various clades becoming more specialized for herbivory over time, perhaps resulting in variation across the primate tree in the ability to exploit insects as a primary dietary resource.

If these constraints, or others, are operating in primates, then they should manifest at the macroevolutionary level as heterogeneity among lineages in the rate of trophic evolution. For example, the hypothesis that diurnality limits the evolution of insectivory predicts that small-bodied anthropoids, which are mostly diurnal, will have a lower rate of transition between trophic states than other small-bodied primate lineages, which are mostly nocturnal. Evolutionary rates have been used to test hypotheses of constraint or to make a posteriori inferences of constraint in a diverse set of organismal traits, including flower size in plants (Barkman et al., 2008), forelimb morphology in marsupials (Cooper & Steppan, 2010), niche evolution in damselfishes (Litsios et al., 2012), growth form in angiosperms (Beaulieu, O’Meara, & Donoghue, 2013), and habitat shifts in diatoms (Nakov, Beaulieu, & Alverson, 2019). The goal of the present study is to evaluate the idea that trophic evolution is constrained in some small-bodied primate lineages by testing for variation in transition rates between insectivory and herbivory against the null hypothesis that a single rate can explain the distribution of insectivory and herbivory across the primate tree.

## 2 MATERIALS AND METHODS

### 2.1 Tree and sample

The analyses reported here were conducted using the phylogenetic topology estimated by Springer et al. (2012) for 367 extant primate taxa. This tree was pruned so that only species-level taxa recognized by Groves (2005) were included, resulting in a tree with 307 tips. Springer et al. provided four sets of divergence dates for the tree based on different assumptions about variation in rates of molecular evolution among lineages and the certainty of fossil calibrations. Two of the timetrees were used for the present study: one that assumed autocorrelated rates of molecular evolution with soft-bounded constraints on fossil calibrations, and one that assumed autocorrelated rates but with hard-bounded constraints. These two trees were preferred over the two that assumed independent rates of molecular evolution because autocorrelated rates provide a much better fit to the primate molecular data and appear to be more biologically realistic (dos Reis et al., 2018). The trees are available in the Supporting Information (Text S1 and Text S2).

Species were classified as insectivorous or herbivorous using primary field reports or recent reviews that compiled information on dietary composition from such reports. A species was considered insectivorous when insects and other small fauna constituted at least 50% of its diet. For some species, dietary percentages were not available. In those cases, assignments were made using qualitative descriptions from experts as long as the characterizations were compatible with quantitative data for the species’ closest living relatives. A total of 26 species were identified as insectivorous (Table 1). The remaining taxa were categorized as herbivorous, which subsumes frugivory, seed predation, folivory, exudativory, and graminivory (see Table S1 in the Supporting Information for the full list of taxa and character coding).

**TABLE 1.** Primates classified as insectivorous for this study

| Species | % Faunivory | Source |
| --- | --- | --- |
| *Galagoides thomasi* | 70 | Nekaris & Bearder, 2007 |
| *Galagoides demidoff* | 70 | Nekaris & Bearder, 2007 |
| *Galago matschiei* | qualitative<sup>†</sup> | Nash, Bearder, & Olson, 1989 |
| *Galago moholi* | 52 | Nekaris & Bearder, 2007 |
| *Galago gallarum* | qualitative<sup>†</sup> | Butynski & de Jong, 2004 |
| *Galago senegalensis* | 50 | Burrows & Nash, 2010 |
| *Paragalago orinus* | qualitative<sup>†</sup> | Rovero, Marshall, Jones, & Perkin, 2009 |
| *Paragalago granti* | qualitative<sup>†</sup> | Génin et al., 2016 |
| *Paragalago zanzibaricus* | 70 | Harcourt & Nash, 1986 |
| *Otolemur garnettii* | 50 | Harcourt & Nash, 1986 |
| *Loris lydekkerianus* | 96 | Nekaris & Bearder, 2007 |
| *Loris tardigradus* | 100 | Nekaris & Bearder, 2007 |
| *Arctocebus calabarensis* | 85 | Rothman et al., 2014 |
| *Arctocebus aureus* | 85 | Rothman et al., 2014 |
| *Allocebus trichotis* | 70 | Biebouw, 2013 |
| *Mirza coquereli* | >50 | Hladik, Charles-Dominique, & Petter, 1980 |
| *Microcebus rufus* | 54 | Rothman et al., 2014 |
| *Tarsius dentatus* | 100 | Niemitz, 1984 |
| *Tarsius tarsier* | 100 | Niemitz, 1984 |
| *Tarsius sangirensis* | 100 | Niemitz, 1984 |
| *Cephalopachus bancanus* | 100 | Niemitz, 1984 |
| *Carlito syrichta* | 100 | Niemitz, 1984 |
| *Saimiri sciureus* | 79–97 | Zimbler-DeLorenzo & Stone, 2011 |
| *Saimiri oerstedii* | 90 | Zimbler-DeLorenzo & Stone, 2011 |
| *Saimiri boliviensis* | 75 | Zimbler-DeLorenzo & Stone, 2011 |
| *Saimiri ustus* | as for other *Saimiri* | Zimbler-DeLorenzo & Stone, 2011 |

<sup>†</sup> For some species, dietary percentages were not available. In such cases, I used qualitative accounts from experts as long as the description was consistent with quantitative data for the species' closest living relatives. See Table S1 in the Supporting Information for the full sample.

Species were further divided into small-bodied and large-bodied using literature compilations of body mass (Smith and Jungers, 1997; Jones et al., 2009; Fleagle, 2013). Two sets of analyses were performed using different size thresholds to evaluate whether the value used to dichotomize body size affects interpretations: 800 g and 1 kg. All species were assigned to a size category based on female body mass, given that females are considered more sensitive to energetic constraints than males (e.g., Gordon, Johnson, & Louis, 2013). Taxa without data on body mass were assigned to a size category when their position relative to the threshold could be assumed with high confidence (e.g., all callitrichines are smaller than 800 g).

The size and diet classifications were combined to create a three-state character: small-insectivorous, small-herbivorous, and large-herbivorous. This character was treated as ordered, with direct transitions between small-insectivorous and large-herbivorous prohibited (i.e., SI↔SH↔LH). This coding scheme allowed transitions between trophic states within small-bodied lineages to be isolated in the analysis without compromising phylogenetic sampling by excluding large-bodied lineages.

### 2.2 Models of trait evolution

Transition rates between character states were estimated using the multistate speciation and extinction model (MuSSE) in the package diversitree (FitzJohn, 2012) for R (R Core Team, 2019). The hypothesis of variation in rates of trophic evolution was tested using diversitree’s make.musse.split function, which splits subclades (foreground clades) from the rest of the tree (paraphyletic background) and allows each partition to have separate rate classes. The locations of the splits are selected prior to analysis. Three foreground clades were used for this study: Anthropoidea, Lemuriformes, and Lorisiformes. Initially, models with one split—one foreground clade and the background—were examined. Depending on the results of those analyses, the model set was expanded to include models with two foreground clades.

For each character state, MuSSE estimates up to three parameters: the transition rate out of the state (*q*), the speciation rate for lineages in the state (λ), and the extinction rate for lineages in the state (μ). Thus, a MuSSE model for an ordered three-state character and no splits will have as many as *k* = 10 estimated parameters. Adding a single split will double that number to *k* = 20, which is almost certainly too many for the size of the primate tree. To reduce the parameter set to a more appropriate size, the following constraints were imposed. First, speciation and extinction rates were not allowed to vary by character state or across partitions. Second, transition rates between insectivory and herbivory in small-bodied lineages were set equal to each other within partitions (i.e., *q*_IH_ = *q*_HI_, where *q*_IH_ is the rate from insectivory to herbivory, and *q*_HI_ is the rate from herbivory to insectivory). Previous analysis of this data set using an unpartitioned tree found that the symmetric-rates model for transitions between trophic states provides a better fit to the data than a model that allows rate asymmetry (i.e., *q*_IH_ ≠ *q*_HI_) (Scott, 2019). Preliminary model comparisons using Akaike’s information criterion indicated that the symmetric-rates assumption is also justified within the partitions examined here. For transitions between size classes among herbivores, there is strong support for rate asymmetry, with the rate into the large-bodied state being several times higher than the rate into the small-bodied state across the primate tree (Scott, 2019). Thus, size transition rates were allowed to vary within partitions. These constraints resulted in two-partition models with *k* = 8 parameters: one trophic transition rate for each partition, two size transition rates for each partition, and one speciation rate and one extinction rate for the entire tree. A three-partition model (two foreground clades and the paraphyletic background) has at least one more parameter (*k* = 9) for the second foreground clade’s trophic transition rate and, depending on the results, two additional parameters for that clade’s size transitions (*k* = 11).

### 2.3 Bayesian analysis of rate estimates

Uncertainty in the maximum-likelihood estimate for each transition rate was examined using a Bayesian approach to approximate each parameter’s posterior distribution. This part of the analysis was conducted with Markov chain Monte Carlo (MCMC) using diversitree’s mcmc function (FitzJohn, 2012). Markov chains were generated following the procedures outlined in Johnson, FitzJohn, Smith, Rausher, & Otto (2011) and FitzJohn (2012), including their use of an exponential prior distribution with a mean of twice the net diversification rate (i.e., speciation rate minus extinction rate) for the entire tree. The chains were run for 120,000 generations, with the first 20,000 being discarded as burn-in. The remaining generations were thinned by sampling every tenth generation, resulting in a final sample of 10,000 generations for further analysis. The R package coda (Plummer, Best, Cowles, & Vines, 2006) was used to examine MCMC diagnostics on the thinned chains. Effective sample sizes for transition rates were high (*n* > 8200, typically *n* > 9000), autocorrelation among generations for each parameter was low (*r* < 0.07, typically *r* < 0.03), and trace plots indicated convergence.

The posterior distributions for the parameter estimates were used to compute posterior probabilities for differences between transition rates. The posterior probability that *q*_*i*_ is greater than *q*_*j*_ is simply the proportion of MCMC samples for which that statement is true (Goldberg et al., 2010). Such comparisons were made across partitions (e.g., Anthropoidea versus the background) and within partitions in the case of size transitions (e.g., the rate of transition into the large-bodied state versus the rate into the small-bodied state in Anthropoidea). Because the posterior distributions for Lorisiformes were strongly right-skewed, rates were log-transformed (base *e*) for visual presentation, but all quantitative comparisons were made using the untransformed rates.

## 3 RESULTS

### 3.1 Two-partition models

With Anthropoidea split from the rest of the tree, there is moderate support for two rates of transition between insectivory and herbivory in small-bodied lineages. Anthropoids have a lower rate than other primates (Figure 2A): the maximum-likelihood estimate for the background rate is approximately 5–7 times higher than the estimate for anthropoids (Table S2). The posterior probabilities for rate heterogeneity in this partitioning scheme range from *PP* = 0.906 to 0.959, depending on the tree and size threshold (Table 2). Support is highest when using a size threshold of 1 kg and the tree with soft-bounded constraints on fossil calibrations.

**TABLE 2.** Support for trophic-rate differences in two-partition models

| Model comparison | Posterior probability |  |  |  |
| --- | --- | --- | --- | --- |
|  | AUTOsoft tree |  | AUTOhard tree |  |
|  | t.800 | t.1000 | t.800 | t.1000 |
| Anthropoidea < background | 0.917 | 0.959 | 0.906 | 0.946 |
| Lorisiformes > background | 0.985 | 0.992 | 0.983 | 0.989 |
| Lemuriformes > background | 0.622 | 0.676 | 0.614 | 0.647 |

Notation: AUTOsoft = autocorrelated rates of molecular evolution and soft-bounded constraints; AUTOhard = autocorrelated rates of molecular evolution and hard-bounded constraints; t.800 = size threshold of 800 g; t.1000 = size threshold of 1 kg.

**FIGURE 2.**
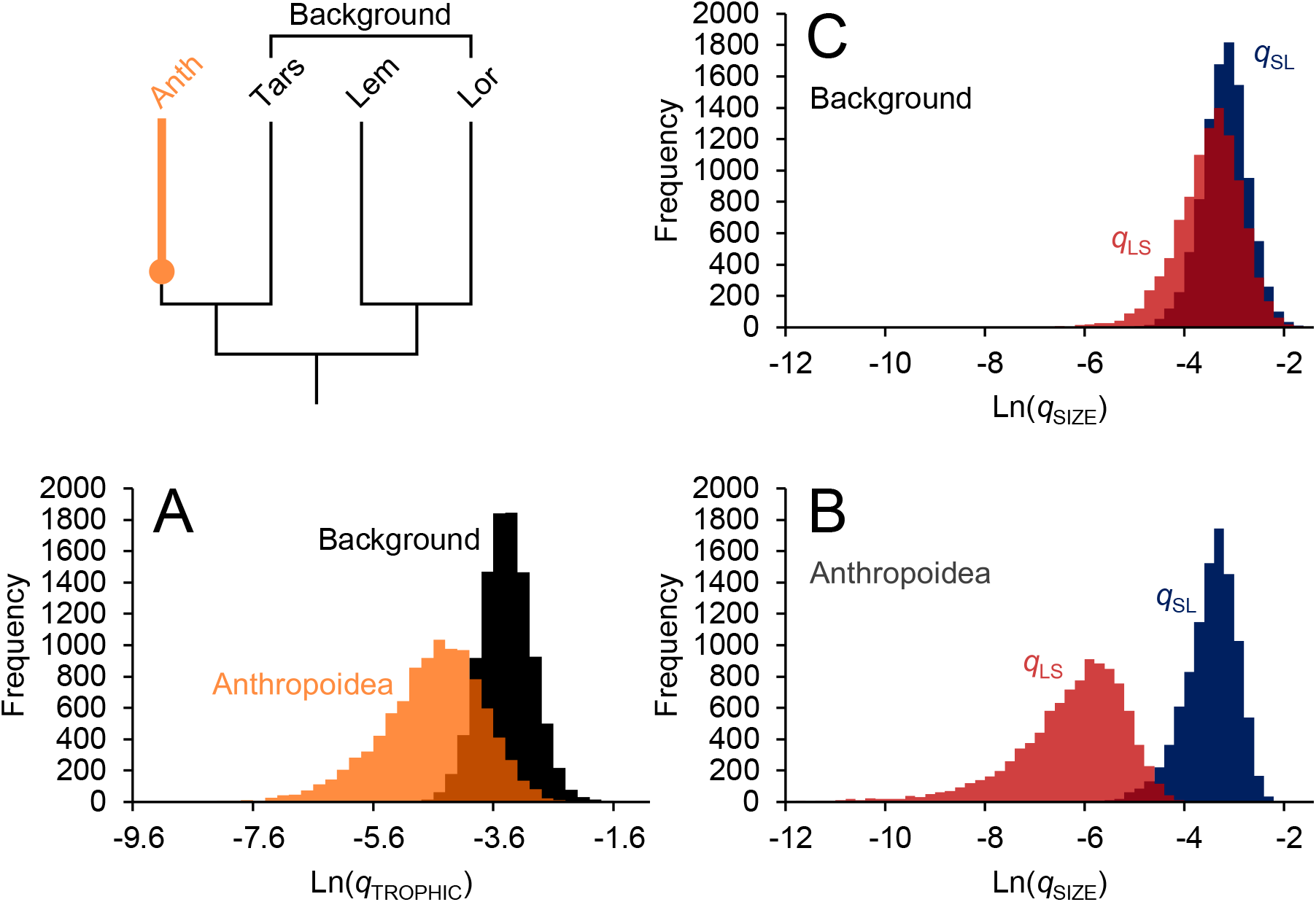
Posterior distributions of trophic transition rates (A) and size transition rates (B, C) for the model with Anthropoidea split from the rest of the tree and a size threshold of 800 g. The results shown here were generated using the timetree that assumed autocorrelated rates and soft-bounded constraints. Anth = Anthropoidea, Tars = Tarsiiformes, Lem = Lemuriformes, Lor = Lorisiformes, *q* = transition rate, SL = small to large, LS = large to small.

There is strong support for asymmetry in size transition rates among anthropoids, where transitions from small to large occur at a much higher rate than transitions in the reverse direction (*PP* > 0.99; Figure 2B). In other primates, there is no evidence for such rate asymmetry (*PP* < 0.70; Figure 2C). This difference in the pattern of size evolution is driven by the very low transition rate out of the large-bodied state in anthropoids. This rate differs from the other size transition rates with high posterior probability (*PP* > 0.98), whereas the other three rates cannot be statistically distinguished from each other (*PP* < 0.85; compare Figure 2B and 2C).

Given the strong support for these contrasting patterns of size evolution, the remaining two-partition models were modified to allow anthropoids to have their own set of size transition rates while constraining the rest of the tree to have a second set of size transition rates, regardless of how the tree was partitioned for the analysis of trophic transition rates. Thus, these models have two partitions for trophic transition rates and two partitions for size transition rates. The model that allowed Lorisiformes to have a distinct trophic rate produced the strongest support for trophic rate heterogeneity among the two-partition models (Figure 3). In this case, lorisiforms have a rate of trophic evolution that is approximately 4–6 times higher than the background rate with high posterior probability (*PP* > 0.98; Table S3). The results from the model that allowed Lemuriformes to have a distinct trophic rate indicate no support for trophic rate heterogeneity (*PP* < 0.70; Figure 4; Table S4).

**FIGURE 3.**
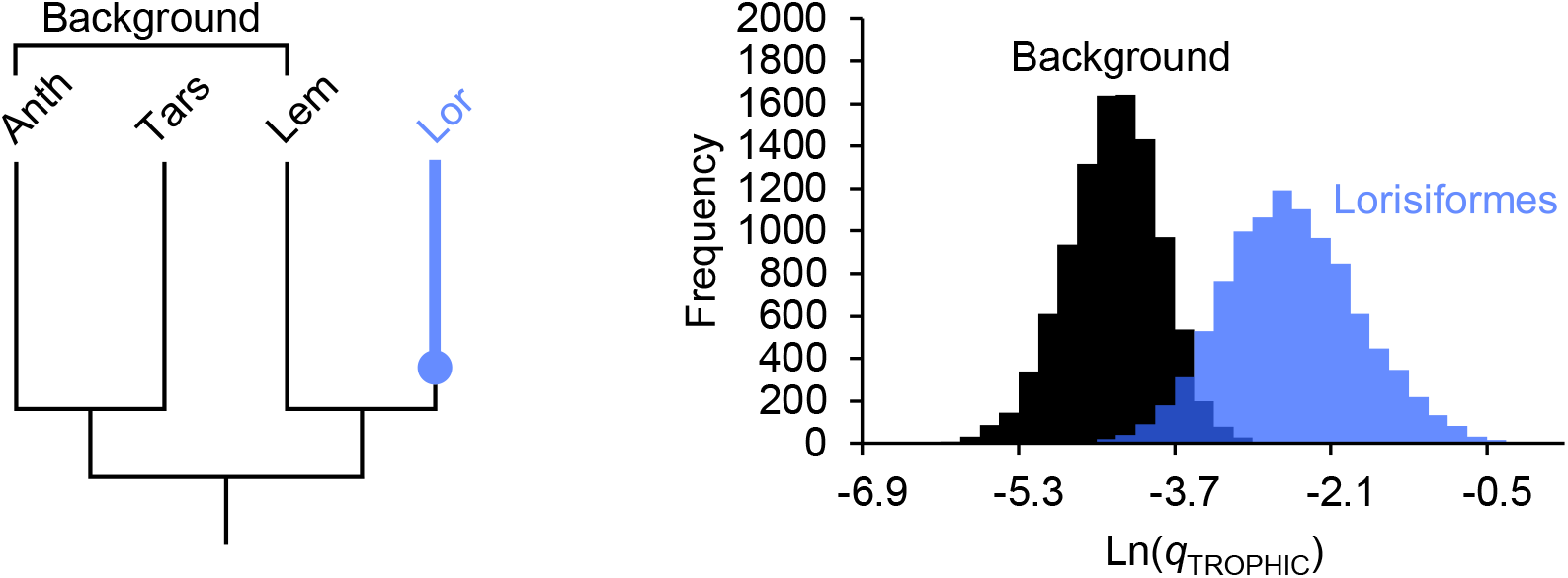
Posterior distributions of trophic transition rates for the model with Lorisiformes split from the rest of the tree and a size threshold of 800 g. The results shown here were generated using the timetree that assumed autocorrelated rates and soft-bounded constraints. Anth = Anthropoidea, Tars = Tarsiiformes, Lem = Lemuriformes, Lor = Lorisiformes, *q* = transition rate.

**FIGURE 4.**
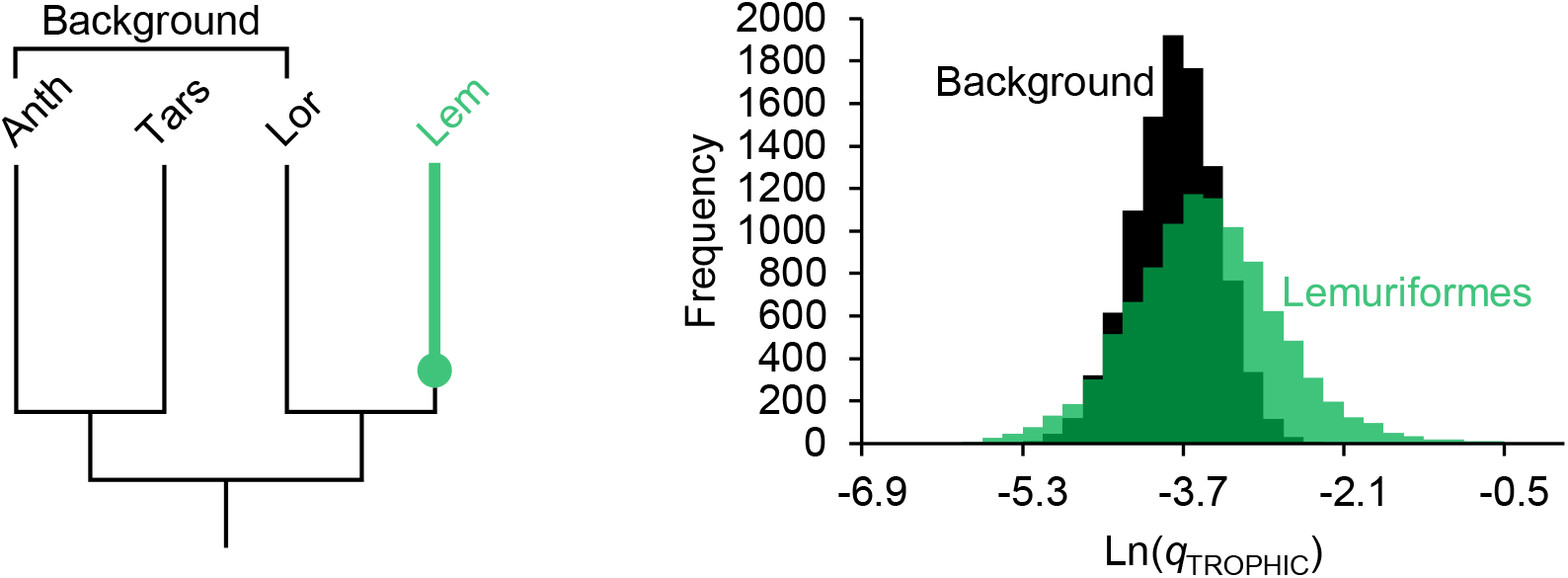
Posterior distributions of trophic transition rates for the model with Lemuriformes split from the rest of the tree and a size threshold of 800 g. The results shown here were generated using the timetree that assumed autocorrelated rates and soft-bounded constraints. Anth = Anthropoidea, Tars = Tarsiiformes, Lem = Lemuriformes, Lor = Lorisiformes, *q* = transition rate.

### 3.2 Three-partition model

The two-partition analyses suggest three possibilities: (1) that anthropoids have a lower trophic transition rate than other primates, (2) that lorisiforms have a higher rate than other primates, or (3) that the two-partition models do not adequately describe the degree of rate heterogeneity in primates. To distinguish among these alternatives, a three-partition model with anthropoids and lorisiforms both foregrounded was constructed. This model allowed each partition to have its own rate of trophic evolution, with size transition rates partitioned as above (i.e., anthropoids versus all other primates, including lorisiforms; Table S5). Analysis of this model indicates that the anthropoid trophic rate cannot be clearly distinguished from the background rate (*PP* < 0.90), that there is moderate support for lorisiforms having a higher trophic rate than the background (0.90 < *PP* < 0.96), and that anthropoids and lorisiforms are very unlikely to be characterized by a common trophic rate (*PP* > 0.98) (Table 3; Figure 5). These results suggest that, of the models considered here, the two-partition model with Lorisiformes as the foreground clade provides the best description of primate trophic evolution.

**TABLE 3.** Support for trophic-rate differences in the three-partition model

| Model comparison | Posterior probability |  |  |  |
| --- | --- | --- | --- | --- |
|  | AUTOsoft tree |  | AUTOhard tree |  |
|  | t.800 | t.1000 | t.800 | t.1000 |
| Anthropoidea < background | 0.791 | 0.861 | 0.787 | 0.838 |
| Lorisiformes > background | 0.939 | 0.951 | 0.933 | 0.950 |
| Lorisiformes > Anthropoidea | 0.986 | 0.995 | 0.984 | 0.991 |

Notation: AUTOsoft = autocorrelated rates of molecular evolution and soft-bounded constraints; AUTOhard = autocorrelated rates of molecular evolution and hard-bounded constraints; t.800 = size threshold of 800 g; t.1000 = size threshold of 1 kg.

**FIGURE 5.**
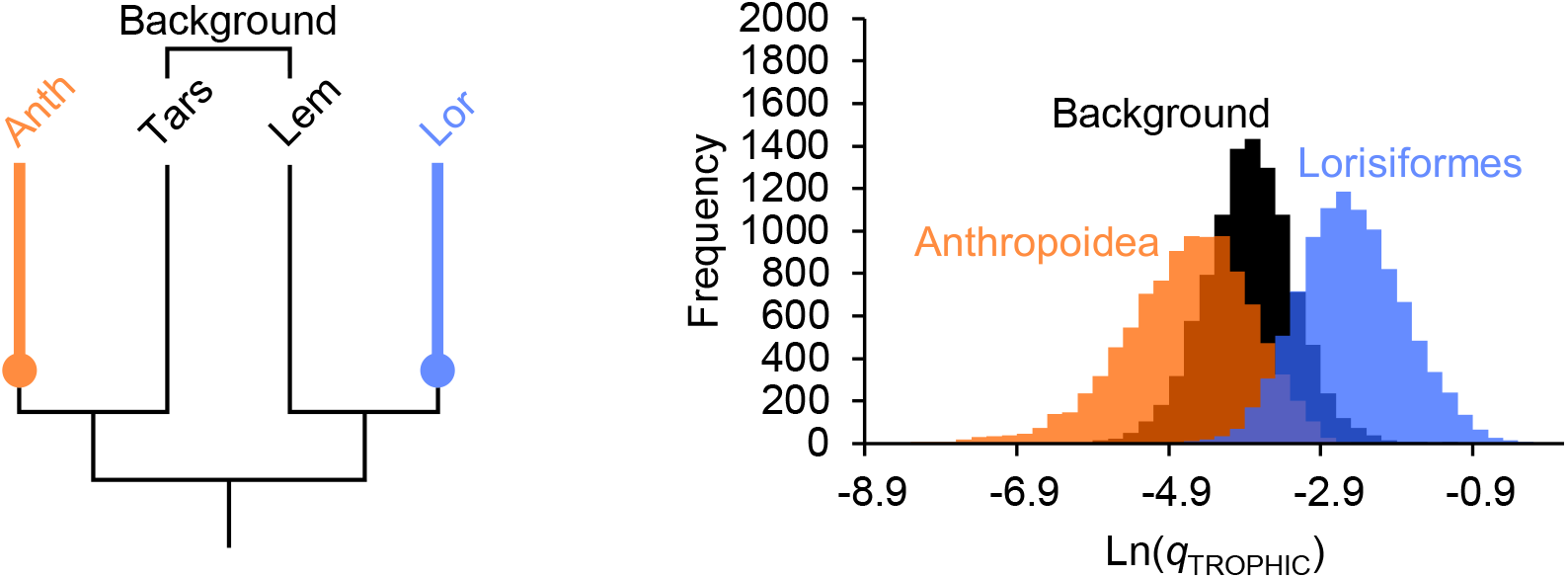
Posterior distributions of trophic transition rates for the model with Anthropoidea and Lorisiformes each split from the rest of the tree and a size threshold of 800 g. The results shown here were generated using the timetree that assumed autocorrelated rates and soft-bounded constraints. Anth = Anthropoidea, Tars = Tarsiiformes, Lem = Lemuriformes, Lor = Lorisiformes, *q* = transition rate.

## 4 DISCUSSION

### 4.1 Activity pattern and trophic evolution

The rate heterogeneity detected here supports the idea that trophic evolution has been more labile in some small-bodied primate lineages than in others. The rarity of primarily insectivorous anthropoids has focused attention on this clade and the possibility that some aspect of diurnal ecology is a constraint on trophic evolution in primates (Cartmill, 1980; Charles-Dominique, 1975; Ross, 1996; Scott, 2019). The results of the present study challenge this idea by showing that a single rate of transition between insectivory and herbivory can explain the distribution of trophic strategies among small-bodied lineages of mostly diurnal anthropoids and nocturnal lemurs and tarsiers. Insectivorous anthropoids and lemurs are nested deeply among herbivorous lineages, indicating that insectivory is an evolutionarily recent phenomenon in these two clades (Figure 1). Thus, despite differences in activity pattern, extant anthropoids and lemurs appear to be characterized by similar trophic evolutionary histories where herbivory has predominated and shifts to insectivory have been infrequent.

What distinguishes anthropoids in this analysis is their pattern of size evolution. In contrast to other primates, which are characterized by symmetric rates of transition into and out of the large-bodied state, anthropoids exhibit high rate asymmetry favoring shifts into the large-bodied state. This result is not surprising in light of the well-known differences in the distribution of body size among primate clades (e.g., Charles-Dominique, 1975; Fleagle, 1978, 2013; Jungers, 1984). The prevalence of large body size (>1 kg) in anthropoids is thought to be one of the solutions to the problem of trophic competition with diurnal birds, allowing anthropoids to specialize on herbivorous resources that birds cannot typically access (e.g., leaves and mechanically protected fruits; Charles-Dominique, 1975; Ross, 1996). The results of this study are consistent with this idea, but they do not constitute additional evidence beyond the observation that anthropoids tend to be larger than diurnal birds. Diurnality can be considered an indirect influence on the distribution of insectivory across the primate tree to the extent that it increases the likelihood that large body size will evolve and persist. This effect is magnified by the tendency of diurnal lineages to diversify and accumulate at a higher rate than nocturnal lineages (Magnuson-Ford & Otto, 2012; Santini, Rojas, & Donati, 2015; Scott, 2018, 2019). However, the results of this study suggest that activity pattern does not have an effect on the rate of transition between herbivory and insectivory among small-bodied primates.

Because the broad-scale phylogenetic approach adopted here does not address the possibility that lineage-specific factors have produced similar patterns of trophic evolution in diurnal anthropoids and nocturnal lemurs, these results should not be viewed as a decisive rejection of the idea that diurnality is a constraint on primate trophic evolution. Studies conducted at a much finer scale of resolution may reveal different processes operating in each clade and establish equifinality. However, given the current state of knowledge, the low frequency of insectivory among small-bodied diurnal anthropoids does not appear to be unusual and therefore in need of explanation. Instead, it is the lorises and galagos that stand out relative to other small-bodied primates in having a greater tendency to shift between trophic states.

### 4.2 Commitment to herbivory as a constraint on the evolution of insectivory

The pattern of rate heterogeneity found in primates is consistent with the idea that adaptive commitment to herbivory has reduced the likelihood that insectivory will evolve or persist in some lineages. There are two ways to interpret the pattern of rate heterogeneity in this context. The first posits that the low rate of transition between trophic states found in most of the primate tree is plesiomorphic, meaning that trophic evolution has been conservative for much of the clade’s history. This inference, combined with the prevalence of herbivory among extant lineages, aligns with the view that many of the apomorphies that unite primates originated as adaptations for acquiring angiosperm products, and that this aspect of the clade’s evolutionary history has biased primates against adopting insectivory as a primary trophic strategy (Rosenberger, 2013; Sussman et al., 2013). It follows that the higher rate of trophic evolution found in lorisiforms represents a derived acceleration, suggesting that lineages in this group evolved traits that allowed them to shift between trophic states more easily than other primates in response to ecological conditions. The evolutionary importance of insects as a primary or secondary dietary resource among lorisiforms was emphasized by Rasmussen & Nekaris (1998), who argued that adaptive divergence between Lorisidae and Galagidae in aspects of locomotor behavior, sensory systems, and life history was driven, in part, by specialization on insects with different properties: cryptic or toxic prey in the case of lorisids versus active and elusive prey in the case of galagids. Notably, the ability to exploit insects has not necessarily channeled lorisiform lineages toward obligate insectivory, as in tarsiers. The present-day expression of this evolutionary history is the gradient of trophic strategies exhibited by galagids and the presence of herbivorous and insectivorous lorisid sister lineages found in both Africa and Asia (Nekaris and Bearder, 2007).

The second scenario posits that the transition rate found in lorisiforms is plesiomorphic and that other primate clades evolved slower rates in parallel as they became more committed to a particular trophic strategy: herbivory in Anthropoidea and Lemuriformes, and insectivory in tarsiers. This scenario is less parsimonious than the first, but there are two lines of evidence that suggest convergent, herbivory-driven rate slowdowns in anthropoids and lemurs. First, as noted above, studies of chitinase genes indicate pervasive homoplasy in loss of function in these genes across primates (Emerling et al., 2018; Janiak et al., 2018). Emerling et al. (2018) inferred that the plesiomorphic number of *CHIA*s for placental mammals is five functional genes, and that tarsiers retain this condition, implying that the last common ancestors of Primates and Haplorhini also had five. The anthropoids and lemurs that have been characterized so far, including small-bodied species, have two or fewer functional *CHIA*s, and some large-bodied species in both clades have lost function in all five genes, indicating separate histories of increasing commitment to herbivory (Emerling et al., 2018; Janiak et al., 2018). This conclusion is further reinforced by the observation that the lorisiform *Otolemur garnettii* has three functional *CHIA*s (Emerling et al., 2018; Janiak et al., 2018).

The second line of evidence suggesting convergent rate slowdowns in Anthropoidea and Lemuriformes is the history of body-size evolution in each clade’s smallest-bodied lineages. The smallest anthropoids are the Callitrichinae, which have long been regarded as phyletic dwarfs (e.g., Ford, 1980; Leutenegger, 1980; Rosenberger, 1992), descended from a common ancestor shared with other platyrrhines that weighed approximately 1–2 kg (Ford & Davis, 1992; Montgomery & Mundy, 2013; Silvestro et al., 2019). The closely related and slightly larger squirrel monkeys (genus *Saimiri*) may also be dwarfed (Ford & Davis, 1992; Rosenberger, 1992; Silvestro et al., 2019). Recent studies of size evolution in lemurs have concluded that the smallest members of this clade—species of the family Cheirogaleidae—have experienced episodes of size reduction similar to those reconstructed for callitrichines (Masters, Génin, Silvestro, Lister, & DelPero, 2014; Montgomery & Mundy, 2013). If these inferences of phyletic dwarfism are correct, then the evolutionary histories of small-bodied anthropoids and lemurs may include long periods of relaxed selection on traits involved in extracting nutrition from insects (e.g., chitinase genes) owing to the lesser importance of insects as a dietary resource at large body size.

That most small-bodied anthropoids and lemurs have apparently entered their current size range via phyletic dwarfism contrasts with the pattern evident in lorisiforms and tarsiers, where small size appears to have prevailed throughout their histories (Beard, 1998; Beard, Qi, Dawson, Wang, & Li, 1994; Jaeger et al., 2010; Rossie, Ni, & Beard, 2006; Steiper & Seiffert, 2012; Seiffert, Simons, & Attia, 2003; Seiffert, Simons, Ryan, & Attia, 2005). The observation that tarsiers and at least some lorisiforms retain more functional *CHIA*s than other primates also suggests long histories of small body size with selection to maintain some of the primitive digestive machinery assembled in early insectivorous mammals (Emerling et al., 2018; Janiak et al., 2018). *Otolemur garnettii* is the only lorisiform in which *CHIA*s have been investigated so far (Emerling et al., 2018; Janiak et al., 2018). It is unclear how typical this galagid is of other lorisiforms, but the fact that the number of functional genes retained by *O. garnettii* (three) is intermediate between tarsiers (five) and anthropoids and lemurs (two or fewer) is consistent with the idea that lorisiforms have experienced episodes of adaptation to herbivory without becoming too specialized, resulting in a clade that has been more flexible than crown anthropoids, lemuriforms, and tarsiers with regard to shifting between trophic strategies.

Thus, whereas the first scenario outlined above views a slow rate of trophic evolution and commitment to herbivory as evolutionarily ancient and tied to the origin of crown primates, the second scenario raises the possibility that trophic evolution in early crown primates was more labile—similar to lorisiforms—before herbivory came to predominate in the case of crown anthropoids and lemurs, and insectivory in the case of tarsiers. Such trophic flexibility is compatible with a last common ancestor of crown primates that was either primarily herbivorous (Rosenberger, 2013; Sussman et al., 2013) or primarily insectivorous (Cartmill, 1974, 1992, 2012), and it implies that the ancestor’s feeding adaptations did not necessarily constrain or bias trophic evolution as the crown clade began to diversify in the late Paleocene and early Eocene.

### 4.3 Evidence for trophic lability in the primate fossil record

The primate fossil record provides some evidence that early small-bodied primates had a greater tendency to shift between trophic states than would be inferred from the distribution of states among extant anthropoids, lemurs, and tarsiers of similar size. Most of the small-bodied primates known from the early and middle Eocene are omomyiforms (e.g., Covert, 1986; Fleagle, 1978, 2013; Gilbert, 2005; Gingerich, 1981). Studies that have examined functional aspects of molar form in this group indicate that it was characterized by a level of trophic diversity similar to that found in extant lorisiforms (Strait, 2001). The evolutionary history of this diversity is difficult to reconstruct with confidence, given uncertainties in the relationships among omomyiform lineages (e.g., Morse et al., 2019; Tornow, 2008; Williams, 1994). Mapping inferred diets onto the phylogenetic tree generated by Seiffert et al. (2018) indicates a minimum of 5–7 shifts between trophic states over the course of approximately 20 million years (Figure 6). By comparison, the minimum number of shifts required to explain the distribution of states among extant primates is 10 across 60 million years or more of evolution (Scott, 2019), suggesting a relatively high rate of trophic evolution in omomyiforms. This conclusion is also supported by evidence for trophic diversity within two of the earliest genera—*Teilhardina* and *Steinius* (Figure 6; Ni et al., 2004; Strait, 2001; Williams & Covert, 1994).

**FIGURE 6.**
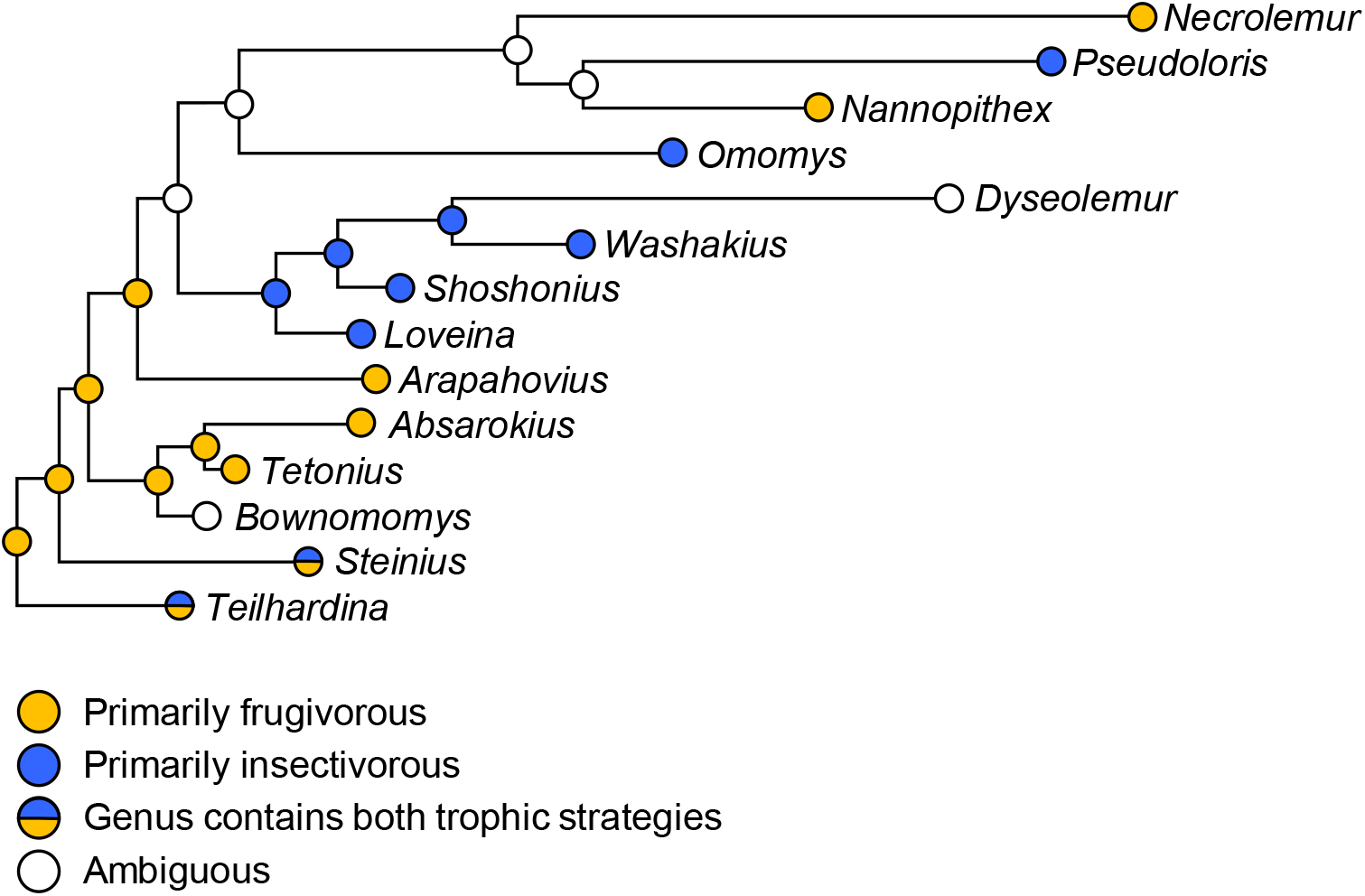
Phylogenetic distribution of trophic strategies among early and middle Eocene omomyiform genera. Dietary reconstructions for each genus are based mainly on the work of Strait (2001), with additional information from Ni, Wang, Hu, & Li (2004) and Williams & Covert (1994). Parsimony reconstructions of diet, obtained using Mesquite (v. 3.61; Maddison & Maddison, 2019), indicate a minimum of 5–7 shifts between trophic states, depending on how ambiguous taxa are coded. The reconstructions shown here are based on the data set where *Bownomomys* and *Dyseolemur* were coded as ambiguous (see Strait, 2001). The tree was taken from the Bayesian tip-dating phylogenetic analysis conducted by Seiffert et al. (2018; see their figure 17). Branch lengths are proportional to time; the tree spans approximately 20 million years from root to most recent tip (*Necrolemur*). *Teilhardina* includes *T. asiatica* and *T. belgica*; *Steinius* includes *S. vespertinus* and *S. annectens*; the use of *Bownomomys* here follows Morse et al. (2019) and is equivalent to *Teilhardina americana* in previous studies. See Strait (2001) and Seiffert et al. (2018) for the complete lists of species-level taxa.

The broader significance of trophic evolution in omomyiforms is unclear owing to a lack of consensus regarding their phylogenetic relationships to the crown clades. Omomyiforms have been interpreted as stem tarsiers, stem haplorhines, or stem primates (see reviews in Fleagle, 2013; Martin, 1993; Miller, Gunnell, & Martin, 2005). Rosenberger (2013), adopting the first of these alternatives as a working hypothesis, argued that trophic diversity within omomyiforms reflects different stages in a shift from frugivory to the highly specialized form of insectivory found in extant tarsiers. According to this view, the implications of omomyiform trophic diversity are limited to the tarsier lineage, and the pattern of diversity mostly indicates a directional macroevolutionary trend of increasing insectivory and its attendant morphological adaptations. However, if omomyiforms are stem haplorhines (e.g., Kay, Ross, & Williams, 1997) or representatives of an early radiation of primates not uniquely related to any of the crown clades (e.g., Martin, 1993; Miller et al., 2005), then their pattern of trophic diversity can be plausibly interpreted as consistent with the hypothesis that trophic evolution in early primates was more labile in comparison to crown Anthropoidea, Tarsiidae, and Lemuriformes.

Other groups of Eocene primates appear to have been less trophically diverse than omomyiforms and more specialized for herbivory. Adapiformes—the other major radiation of primates known from the early and middle Eocene—are mostly large-bodied and are thought to have filled the ecological niches that are now dominated by extant large-bodied anthropoids and lemurs (i.e., diurnal herbivores) (Covert, 1986; Fleagle, 1978, 2013; Gilbert, 2005). Nevertheless, there is some evidence for trophic diversity among early small-bodied members of this group (e.g., *Donrussellia*, *Asiadapis*, and *Marcgodinotius*; Bajpai et al., 2008; Gilbert, 2005). A similar pattern may hold in anthropoids, especially if Eosimiidae are stem anthropoids (Beard, Qi, Dawson, Wang, & Li, 1994; Kay et al., 1997; Williams, Kay, & Kirk, 2010; but see Miller et al., 2005). Small-bodied anthropoids from the late Eocene and early Oligocene have been reconstructed as primarily frugivorous (Kirk & Simons, 2001). In contrast, the molars of middle Eocene eosimiids exhibit morphologies suggesting that these species were more insectivorous than later anthropoids (cf. Heesy & Ross, 2004; Kirk & Simons, 2001) and perhaps comparable to *Saimiri*, the most insectivorous extant anthropoid (Zimbler-DeLorenzo & Stone, 2011; Table 1). Thus, although adapiforms and early anthropoids appear to have been largely herbivorous radiations, there are hints of greater trophic diversity in the earliest members of these clades, suggesting that trophic evolution may have been more labile before herbivory became the dominant trophic strategy.

## 5 CONCLUSIONS

The results of this study indicate that the rate of trophic evolution in small-bodied primates varies among clades. Contrary to expectations, small-bodied anthropoids do not have an unusually low rate in comparison to other lineages. This finding challenges the hypothesis that there is a direct connection between diurnality and the low frequency of insectivorous anthropoids. The main contrast detected here involves lorisiforms, which have a much greater tendency to shift between insectivory and herbivory than other primates. The implications of this pattern of rate heterogeneity are unclear. The most parsimonious interpretation is that the lorisiform rate is apomorphic, implying that primate trophic evolution has been conservative throughout much of the clade’s history. However, various lines of evidence suggest the possibility of convergent rate slowdowns in anthropoids, lemuriforms, and tarsiers owing to greater specialization for herbivory in the case of anthropoids and lemurs, and insectivory in the case of tarsiers. These two scenarios can be tested as sampling of the earliest part of the primate fossil record increases, and as our understanding of the Eocene primate phylogeny and trophic adaptations improves.

## ACKNOWLEDGMENTS

I thank Thierra Nalley and Kristi Lewton for helpful comments on an earlier version of this manuscript.

## DATA AVAILABILITY STATEMENT

The data used for this study are available in Table S1 in the Supporting Information.

